# Robust planar morphogenesis in leaf-like organs requires callose-dependent coordination of division and growth

**DOI:** 10.64898/2026.11.05.764497

**Authors:** Wenye Lin, Stéphanie Hallet, Petra Cifrova, Mandar Deshpande, Daniel Grégoire, Alexis Lacroix, Roman Pleskot, Richard S. Smith, Daniel Kierzkowski, Yoan Coudert

**Author notes:** denotes equal contribution (Stéphanie Hallet, Petra Cifrova).

## Abstract

Planar organs, such as leaves, are key developmental innovations underlying the evolutionary success of plants. In vascular plants, building a planar organ requires coordinating cell division and growth both within the organ plane and across multiple, mechanically distinct cell layers. Phyllids, the leaf-like organs of bryophytes, typically consist of a single cell layer and offer a minimal system to decouple in-plane growth coordination from inter-layer mechanics. Here, we investigate how robust planar morphogenesis is achieved in phyllids, using the moss *Physcomitrium patens* as a bryophyte model. We show that impaired cytokinesis, caused by the loss of a callose synthase gene, disrupts growth gradients, ultimately yielding smaller, non-planar organs of variable shape. Combining quantitative live imaging with computational modeling, we show that the severity of shape defects in mutants depends on the spatial distribution and developmental timing of local growth arrests. These findings reveal how callose-dependent coordination between cell division and in-plane growth prevents mechanical conflicts and maintains planarity in a monolayered plant organ.

## Introduction

Plant organs acquire their diverse three-dimensional shapes through differential growth, *i.e.,* spatial and temporal variations in expansion rate, anisotropy, and direction across cells and tissues (1, 2). Because plant cells are encased within a rigid cell wall and cannot migrate, growth differences between cells and tissues can generate mechanical conflicts (3, 4). These growth-derived conflicts can be critical drivers of organ shape development (5–7), but they also need to be locally buffered to maintain planar geometries (8–10). Developing and maintaining a flat organ requires strict coordination of growth at multiple scales: between the adaxial and abaxial sides of the organ, between distinct anatomical regions such as the midrib, petiole, lamina (blade) and margin, and within the plane of the organ itself, where gradients in growth rate or anisotropy can generate compressive stresses sufficient to cause buckling or wrinkling (11–13). Our understanding of how this coordination is achieved has advanced considerably through the study of *Arabidopsis thaliana* leaves and sepals as model systems (9, 14–19). However, these organs are multilayered structures comprising distinct tissue types, each with different mechanical properties, growth rates, and cellular architectures, which makes it difficult to isolate the contribution of individual cellular processes to overall organ planarity.

Phylids, the leaf-like organs of bryophytes, evolved planar shapes independently of vascular plant leaves. In the moss *Physcomitrium patens*, the phyllid lamina is a one-cell-thick layer, which makes cell division and growth directly observable and offers a minimal framework for dissecting planar morphogenesis (20). Despite this major structural difference from vascular plant leaves, recent work in *P. patens* has uncovered strikingly similar morphogenetic principles (21). Phyllids undergo an early proliferative phase driven by a basal zone of cell division, followed by cell expansion as cells exit the division zone. Cells of the developing midrib cease anticlinal divisions and elongation before lamina cells, while spatial and temporal growth gradients across the lamina collectively determine the final organ shape (21). Time-lapse imaging combined with quantitative cellular analyses has established the phytohormone auxin as a major positional cue promoting cell expansion and differentiation (21–24). Despite the identification of various molecular regulators of phyllid development (20, 23, 25–29), the mechanism that coordinates cell division with growth gradients remains unknown.

A direct way to test how cell division contributes to planar morphogenesis is to perturb it and examine the consequences for organ shape. Cytokinesis – the final stage of cell division – ends with the centrifugal expansion of the cell plate until it fuses with the parent wall (30, 31). Callose, a β-1,3-glucan polysaccharide synthesized by callose synthase (CALS) proteins (also called glucan synthase-like, GSL), accumulates transiently at the expanding cell plate and scaffolds the deposition of other cell wall components, so reducing callose synthesis impairs wall completion (32–37). In *A. thaliana*, *cals10/gsl8* mutants fail to complete wall closure and show severe alterations of leaf shape and flatness, but the multilayered structure of the leaf makes it difficult to relate defective cytokinesis in individual cells to altered planarity (38–40). *CALS* genes are conserved across land plants, with 12 putative members in the moss *P. patens*, and their role in cytokinesis likely predates land plants, as suggested in the streptophyte alga *Penium margaritaceum* (41–44). This evolutionary conservation makes CALS-dependent cytokinesis a tractable entry point to interrogate how cell division shapes planar organs. Here, taking advantage of this conserved feature, we investigate how the coordination between cell division and growth drives robust morphogenesis in the moss phyllid.

## Results

### *Physcomitrium patens CALS6* regulates phyllid shape robustness

To explore the role of callose in cell division during phyllid development, we stained dissected *P. patens* gametophore (leafy shoot) apices with aniline blue and imaged the fluorescence in emerging phyllids by confocal microscopy. At the 2-2.5-day stage, the signal coincided with cell walls across the whole organ, whereas at the 3-day stage it was mostly restricted to the proximal region, which corresponds to the zone of active cell division (21). This dynamic pattern indicates that callose is actively deposited and then removed as phyllids grow (Figure 1A-1B).

**Figure 1.**
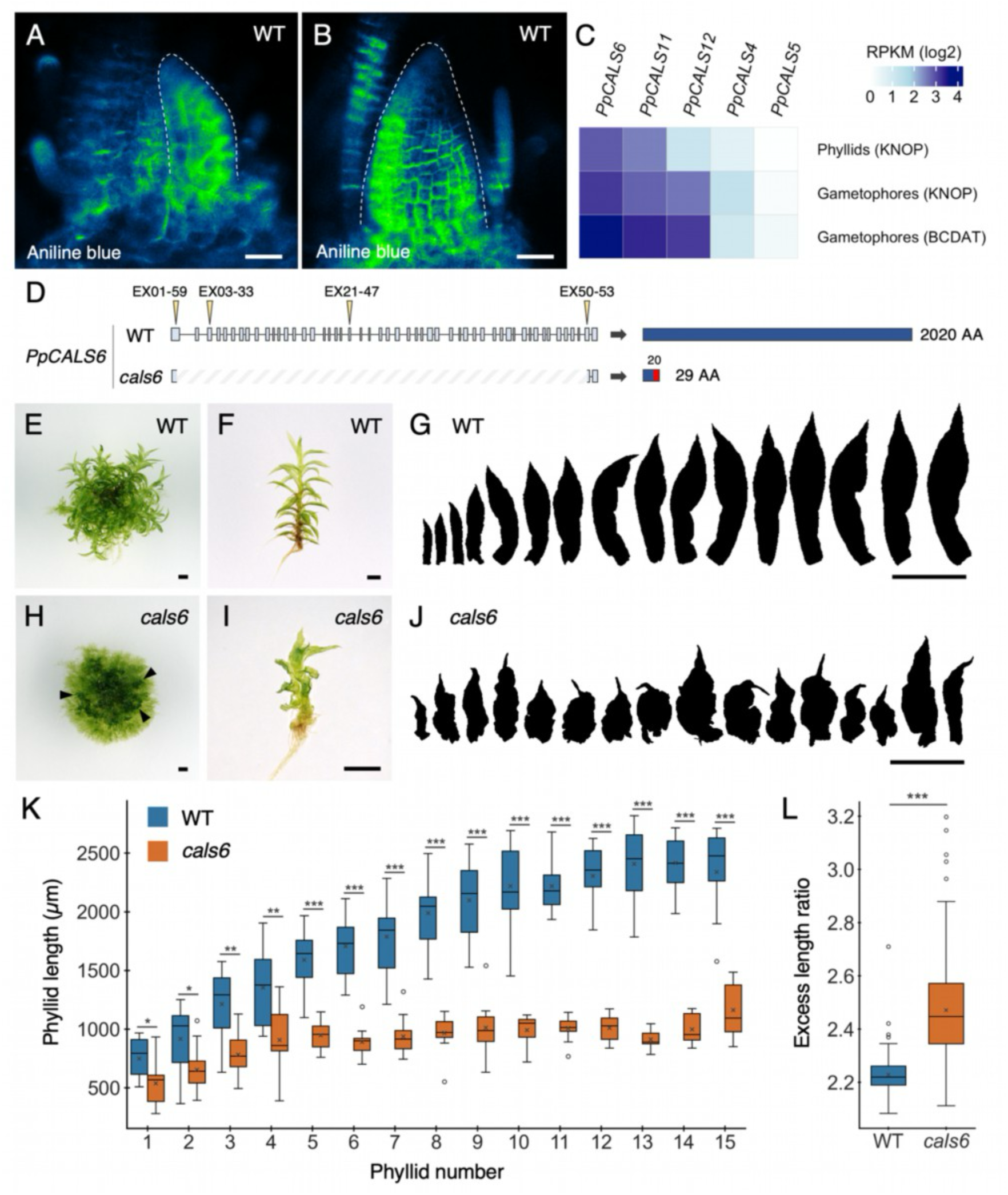
Robust phyllid morphogenesis in *Physcomitrium patens* requires *CALS6*. (**A-B**) Callose deposition patterns revealed by aniline blue fluorescence in emerging phyllids of wild-type (WT) gametophores. Dotted lines mark phyllid outlines. (**C**) Heat map showing that *CALLOSE SYNTHASE6 (CALS6)* is the most highly expressed gene out of five *CALS* genes detected in *Physcomitrium patens* phyllids and gametophores, grown either on KNOP or BCDAT medium. (**D**) Molecular characterization of the *cals6* CRISPR mutant (line #2). Left: diagrams showing the *CALS6* locus in WT and *cals6* mutant backgrounds, with light grey-blue boxes showing exons, yellow arrowheads indicate the positions of sgRNAs, and dashed lines indicate the edited region identified in the *cals6* mutant by Sanger sequencing. Right: schematic representations of the predicted proteins, with WT sequence shown in dark blue and predicted frameshifted regions in the *cals6* mutant shown in red. Numbers indicate predicted amino acid (AA) numbers. (**E-J**) Representative colonies (E, H), gametophores (F, I) and phyllid developmental series (G, J) showing developmental alterations in *cals6* mutants (H-J) compared with WT (E-G). (**K**) Quantification of phyllid length along the gametophore axis, revealing that *cals6* mutants (*n* ≥ 7) have shorter phyllids than WT (*n* ≥ 9). (**L**) Phyllid shape complexity (excess length ratio = phyllid perimeter divided by phyllid length) is greater in *cals6* mutants (*n* = 192) than in WT (*n* = 149). For (K) and (L), statistical significance of a Welch t-test and a Mann-Whitney U one-sided test, respectively, is indicated as follows : *, (0.01 < p ≤ 0.05) ; **, (0.001 < p ≤ 0.01) ; ***, (p ≤ 0.001). Boxes show the interquartile range, the horizontal line the median, and dots the mean; whiskers extend to the most extreme values within 1.5× the interquartile range. Scale bars : A-B, 20 µm ; E-J, 1 mm. For all panels, *CALS6* refer to the *P. patens* gene *PpCALS6*.

Next, we used a genetic approach to alter callose biosynthesis. Among the five *CALS* genes expressed in phyllids and gametophores, *CALS6* had the highest expression levels in all conditions tested (Figure 1C). To examine its function, we disrupted its coding sequence by CRISPR-Cas9 genome editing. Using four sgRNAs, we deleted a chromosome fragment of more than 18 kb spanning nearly the entire *CALS6* coding sequence. The deletion introduced a frameshift at codon 20 and was predicted to yield a truncated protein of 29 amino acids instead of the 2,020 amino acids of the wild-type protein (Figure 1D).

To assess the developmental consequences of loss of *CALS6* function, we compared the gross morphology of *cals6* mutants with that of wild-type controls (Figure 1E, 1H). Gametophores of *cals6* mutants were stunted, with an average length more than 50% shorter than in wild type (Figure 1F, 1I; Figure S1). In wild type, phyllids are heteroblastic: their length increases gradually during the juvenile-to-adult transition and stabilizes around the 10^th^ position from the base, and adult phyllids have a lanceolate shape (Figure 1G, 1K) (22, 45). In *cals6* mutants, phyllids were shorter at all positions, and the heteroblastic gradient was less pronounced, with length reaching a plateau around the 4^th^ position (Figure 1K). Mutant phyllids were also more irregular in shape, with crinkled margins, as reflected by a higher excess length ratio, defined as the phyllid perimeter relative to its length (Figure 1J, 1L) (46). Independent lines generated with the same sgRNAs, but carrying different mutations showed similar phenotypes, supporting the conclusion that loss of *CALS6* function causes these defects (Figure S1). Together, these results identify *CALS6* as an essential regulator of phyllid morphogenesis in *P. patens*.

### *CALS6* regulates tissue planarity in phyllids

To elucidate the cellular basis of the morphological defects of *cals6* mutants, we segmented individual cells in mature adult phyllids using MorphoGraphX and quantified their area and curvature. The proximal–distal gradient of cell area typically observed in wild type was disrupted in *cals6* phyllids, which showed frequent patches of juxtaposed small and large cells (Figure 2A). In addition, whereas wild-type cells were predominantly elongated and aligned along the proximal–distal axis, *cals6* mutant cells were frequently irregular in shape and smaller overall, suggesting impaired growth (Figure 2I). In wild type, cell curvature followed a consistent medio-lateral gradient, with negative values along the midrib, near-zero values across the blade, and positive values along the margins (Figure 2B, 2J). Cross-sections confirmed that the two halves of the blade flanking the midrib were flat at the tissue level, consistent with the distribution of cell curvature values along the proximal-distal axis. In contrast, the medio-lateral curvature gradient was severely impaired in *cals6* mutants (Figure 2B). Although margin cells retained an overall positive curvature similar to wild type, cell curvature across the blade and midrib was irregular, resulting in greater variability in curvature values than in wild type (Figure 2J). Consistent with this, cross-sections of mutant blades revealed a loss of planarity (see insets in Figure 2B).

**Figure 2.**
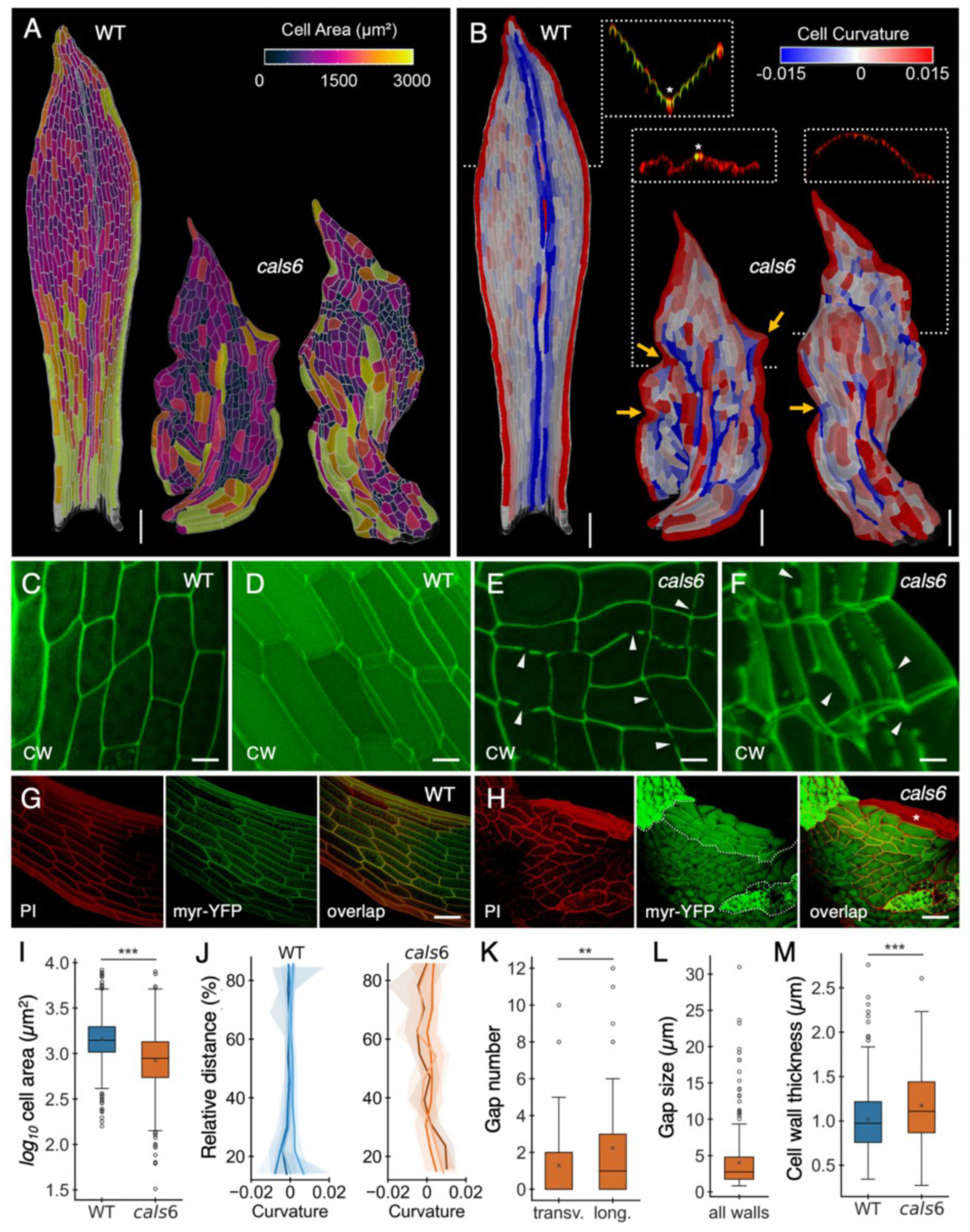
Cellular defects account for shape variability and planarity loss in *cals6* phyllids. (**A-B**) Segmented confocal images of isolated phyllids showing the distribution of cell area (A) and curvature (B) in wild type (WT) and *cals6* mutants. In (B), insets show cross sections through phyllids at the indicated positions, yellow arrows indicate protrusions. (**C-F**) Confocal images of cell wall (CW) autofluorescence in WT and *cals6* mutant phyllid blades, shown as 2D surface views (C, E) and 3D projections of cell morphology (D, F). Arrowheads indicate discontinuities and gaps in cell walls. (**G-H**) Confocal images of phyllid blades in WT (G) and *cals6* mutants (H) expressing a plasma membrane marker (myr-YFP) and stained with propidium iodide (PI), revealing altered membrane organization in *cals6* mutants. Dotted lines delimit putative clonal sectors exhibiting similar alterations. The asterisk marks dead cells. (**I**) Boxplots showing reduced cell area in *cals6* mutants compared with WT (WT, *n* = 1599 ; *cals6*, *n* = 1039). (**J**) Plot showing increased variability in cell curvature in *cals6* compared with WT. Lines show the median curvature within distance bins and shaded bands indicate the interquartile range for *n* independent phyllids (WT, *n* = 3 ; *cals6*, *n* = 4). (**K**) Boxplots showing the number of gaps in transverse (trans.) (n = 88) and longitudinal (long.) (n = 89) cell walls in *cals6* mutants. (**L**) Boxplot showing the distribution of gap sizes in *cals6* mutants (*n* = 348). (**M**) Boxplots showing increased cell wall thickness in *cals6* compared with WT (WT, *n* = 252 ; *cals6*, *n* = 208). For (I, K, M), the statistical significance of a Mann-Whitney U one-sided test is denoted as follows: **, (0.001 < p ≤ 0.01) ; ***, (p ≤ 0.001). Scale bars: A-B, 100 µm ; C-F, 20 µm and G-H, 50 µm. For all panels, *CALS6* refer to the *P. patens* gene *PpCALS6*.

### Cellular architecture is altered in *cals6* phyllids

To determine whether the cellular defects were associated with impaired cytokinesis, we performed confocal imaging of mature phyllid laminae. Two-dimensional surface views revealed frequent cell wall discontinuities in the *cals6* mutant, whereas wild-type cells displayed intact walls throughout (Figure 2C, 2E). Three-dimensional (3D) reconstructions showed that these discontinuities corresponded to gaps of diverse sizes and geometries within the cell walls (Figure 2D, 2F). Quantification of gap number per cell wall in 3D confocal images showed that longitudinal walls contained more gaps than transverse walls, probably because of their larger surface area (Figure 2K). Of all gaps, 73% were located at cell edges, 14% occurred within the wall faces away from the edges, and 13% spanned the entire wall from one edge to the other, creating large openings between adjacent cells. Most gaps were smaller than 5 µm, but some extended beyond 30 µm (Figure 2L). Cell wall thickness was more variable and, on average, greater in *cals6* mutants than in wild type, suggesting that loss of *CALS6* function likely alters cell wall assembly (Figure 2M).

To test whether these defects were accompanied by changes in the plasma membrane, we knocked out *CALS6* by CRISPR/Cas9 in a transgenic line constitutively expressing a YFP fused to an N-terminal myristoylation signal (21). We stained cell walls of the resulting *cals6* mutant with propidium iodide (PI), which also labels dead cells as it penetrates compromised membranes. This dual labelling revealed groups of cells with an altered appearance in the mutant (*e.g.*, loss of YFP signal and PI-positive cytoplasm), which was not seen in wild type and suggested a local loss of membrane integrity (Figure 2G, 2H). Together, these results indicate that *CALS6* is required to maintain cellular architecture, including cell wall and plasma membrane integrity.

### Altered cell division and growth gradients underpin developmental defects in *cals6* phyllids

To investigate how morphological perturbations arise during *cals6* phyllid development, we performed time-lapse imaging of upper phyllids at 0.5-day intervals and compared their growth patterns with those of wild-type, using the protocol we previously established (21). The number and arrangement of merophytes (*i.e.*, groups of cells each derived from a single initial cell; delimited by white lines in Figure 3A-3D) were similar in *cals6* mutants and wild type, suggesting that phyllid initiation, marked by the oblique division of the initial cell, proceeds normally in mutants. Two days after initiation, *cals6* phyllid primordia had a shape and size comparable to wild type (Figure 3A-D). The total duration of phyllid development, approximately 5.5 days in wild type, was also unchanged in *cals6* mutants (Figure 3A-3D). A basipetal growth gradient, in which cells progressively cease dividing and expanding from the phyllid tip toward the base, was detected in both genotypes (Figure 3A-3D). In the mutant, however, the actively growing region was smaller and confined to more proximal positions (Figure 3A-3D). Within this region, fewer cells were dividing or expanding, and the rates of cell division and area expansion were lower than in wild type (Figure 3E-3F), consistent with the smaller cells and reduced cell number in mature *cals6* phyllids (Figure 2A-2I, 3E). The cellular growth gradient within the actively growing zone was also less pronounced in *cals6*, with a more heterogeneous distribution of expanding cells (Figure 3A-3D, 3G-3J). In addition, growth at the base of wild-type phyllids, where they attach to the stem, was strongly inhibited, giving the base a characteristic narrowing (Figure 3C, 3I). This pattern was absent in *cals6*, which probably explains its broader phyllid base (Figure 3D, 3J). Thus, loss of *CALS6* function lowers cell division and growth rates and makes growth more heterogeneous. Because the duration of development is unchanged, this alters the developmental gradients and results in smaller phyllids.

**Figure 3.**
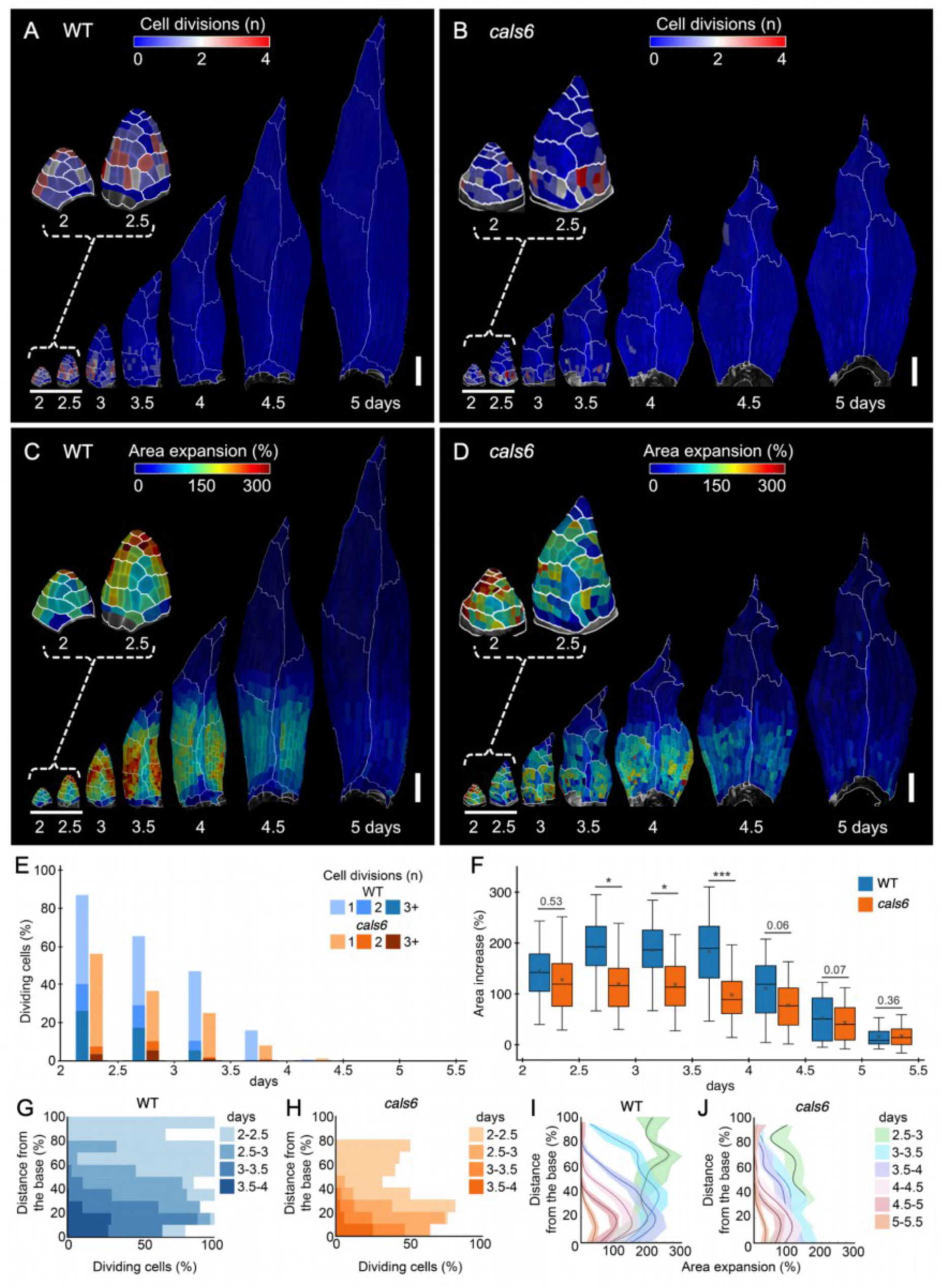
Growth and division gradients driving phyllid morphogenesis are altered in *cals6* mutants. (**A-D**) Heat maps of cell divisions (A-B) and area expansion (C-D) in wild-type (WT) (A,C) and *cals6* (B,D) upper phyllids. Heat values are displayed in the earlier time points. Color scales are similar in (A) and (B), and (C) and (D). (**E-F**) Plots showing that cell division (E) and area expansion (F) are overall reduced in upper *cals6* phyllids compared with WT (n = 928 cells). In (F), Boxes show the interquartile range, horizontal lines the median, and crosses the mean; whiskers extend to the 5th to 95th percentiles. Statistical significance of a Welch t-test is indicated as follows: *, (0.01 < p ≤ 0.05) ; ***, (p ≤ 0.001). (**G-J**) Quantification of cell division (G-H) and area expansion (I-J) as a function of normalized distance from the organ base, showing overall reduced cell division and altered growth gradients in *cals6* mutants (H, J) with respect to WT (G, I). In (I-J), lines correspond to the median, and shaded areas represent the interquartile range. For (E-J), data were computed from 3 and 4 independent WT and *cals6* mutant phyllids, with a maximum number of 928 and 672 cells, respectively. Scale bars: A-D, 100 µm. For all panels, *CALS6* refer to the *P. patens* gene *PpCALS6*.

### Timing and location of local growth arrests account for *cals6* phyllid shape defects

Although altered patterns of cell division and growth were observed in all *cals6* mutant phyllids analyzed, their morphological consequences differed substantially between organs, including within the same gametophore, reflecting a general loss of shape reproducibility (Figure 1J). This variability makes it difficult to dissect the individual contributions of the defects by live imaging alone. It suggests that the defects arise stochastically during development, with both their timing and their location likely influencing their impact on final organ shape. To test whether such defects could account for the abnormal shapes, we used a previously developed computational model, which qualitatively recapitulates the growth dynamics of developing phyllids by using positional cues to control growth and cell division (21). We asked whether the model could reproduce the morphological abnormalities of mutant phyllids by introducing local growth arrests that mimic the cell expansion defects observed in the mutant.

First, we simulated growth arrest in multiple cells randomly but evenly dispersed throughout the lamina (Figure 4A-4B). These perturbations altered local cell organization and reduced organ size, but overall phyllid shape remained largely similar to that of control simulations. The model also predicted that early growth arrests are more detrimental than equivalent arrests occurring later, consistent with cells formed early contributing more to final organ size. These simulations qualitatively reproduced the effects of isolated single-cell growth arrests observed in *cals6* mutants (Figure 4C-4D).

**Figure 4.**
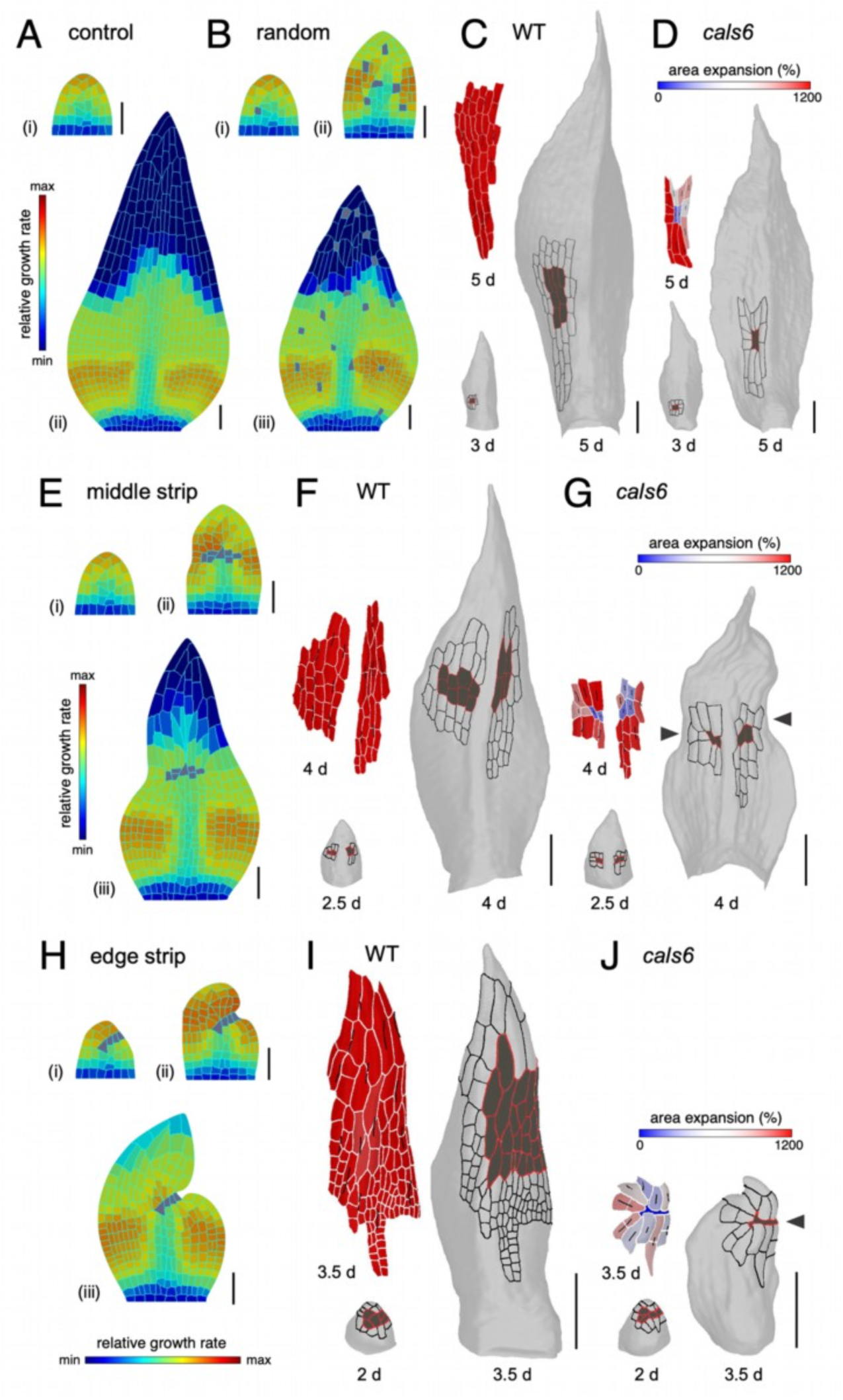
Stochastic local growth inhibition events contribute to global shape defects in *cals6* phyllids. (**A-B, E, H**) Model outputs at successive time points showing phyllid shape in the reference simulation (A) and in simulations with growth arrests imposed on single cells at random positions across the blade (B), or on a strip of cells in the middle (E) or at the edge (H) of the blade. Simulations A(i), B(i), E(i) and H(i) correspond to time frame 120; B(ii), E(ii) and H(ii) to time frame 165; A(ii) and B(iii) to time frame 275; E(iii) to time frame 245; and H(iii) to time frame 215. (**C-D, F-G, I-J**) Heat maps of cumulative area expansion in various wild-type (WT) and *cals6* mutant upper phyllids, from 2.5 to 5 days of growth. Black lines indicate the direction of maximum growth. The heat maps were computed using the parent labels from the early time point and displayed on the later time point. Arrowheads in (G) and (J) indicate protrusions. Scale bars: A-B, E, H = 50 µm; C-D, F-G, I-J = 100 µm. For all panels, *CALS6* refer to the *P. patens* gene *PpCALS6*.

Next, we simulated growth arrest in a group of cells in the central region of the lamina (Figure 4E). This produced pronounced tissue-level defects, including constrictions that became more severe as development progressed, showing that, in the model, coordinated arrest across larger groups of cells has much stronger morphological consequences than dispersed arrests. These simulations resembled the effects of arrested growth in central cell patches observed in *cals6* mutants (Figure 4F-4G).

Finally, when multiple cells located at the margin were prevented from growing, the model produced pronounced indentations (Figure 4H). Marginal growth arrest in *cals6* phyllids also altered shape, but such indentations were not systematically observed. Instead, neighboring cells appeared to compensate partially for the arrest by expanding toward the region resistant to growth, as shown by the orientation of growth tensors at 3.5 d (Figure 4I-4J). This suggests intercellular feedback, possibly mechanical, that buffers local growth defects during tissue growth. Together, these results indicate that local growth arrests can account for the abnormal morphology of *cals6* phyllids, with consequences that depend strongly on their timing and spatial distribution.

## Discussion

In vascular plants, organ planarity depends on a strict coordination of growth rate and orientation both within the organ plane and between cell layers (11). In contrast, moss phyllids offer a minimal system to isolate the in-plane component of this problem, independently of the inter-layer mechanics that dominate most angiosperm models. Here, through a detailed quantitative analysis of phyllid morphogenesis in the *P. patens cals6* mutant, we reveal how the coordination between cell division and growth contributes to the robust shaping of these organs.

Our data reveal that *P. patens CALS6* disruption strongly impairs both cell division and growth processes, producing organs with variable, non-planar shapes. The similarity of these phenotypes to those of *A. thaliana cals10/gsl8* mutants supports the idea that *CALS*-dependent regulation of cell division is evolutionarily conserved across distantly related land plant lineages (38, 40, 41). This result is also consistent with the broader observation that moss phyllids and angiosperm leaves share a common developmental logic. In both systems, tightly controlled spatio-temporal growth heterogeneities across the lamina underlie robust planar shape establishment (11, 20, 21). Using live imaging and computational modeling of the stochastic defects induced by *cals6* loss-of-function, we found that local growth perturbations can produce minor or severe shape defects depending on their timing and spatial location. Because the phyllid lamina consists of a single cell layer, these defects cannot result from a mismatch between mechanically distinct cell layers, as proposed in *A. thaliana* (9, 18, 47). Instead, localized within-layer growth arrests are sufficient to generate growth conflicts that compromise shape robustness. Recent work in *A. thaliana* showed that *jaw-D* mutant leaves curve because proximal–distal growth is uneven across the width of the blade, not because of differential growth between the adaxial and abaxial sides, indicating that in-plane growth coordination is also required for flatness in a multilayered organ (16). Whereas the *A. thaliana jaw-D* mutant reflects a systematic imbalance between midrib and blade, the *P. patens cals6* mutant shows that cell-scale growth arrests are sufficient to produce a comparable outcome in a monolayered organ.

Our observations indicate that some growth defects in *cals6* phyllids are at least partially buffered by the surrounding tissue, possibly through mechanical feedback between neighboring cells. This buffering capacity parallels findings in *A. thaliana*, where robust patterning is achieved despite substantial cell-autonomous stochasticity, and growth conflicts are mechanically compensated to avoid buckling (48, 49). Tolerance to local, random variations in cellular growth may thus be a general requirement for achieving reproducible planar organ shapes across land plants. However, the out-of-plane tissue deformations observed in *P. patens cals6* phyllids reveal that some growth conflicts are resolved at the expense of planarity. Notably, marginal growth arrest was not fully compensated by neighboring cells, consistent with the distinct morphological identity of margin cells relative to lamina cells (21, 45), and the notion that margin domains may act as a structural frame constraining the expanding blade (11, 12). This suggests that this frame-like role of the margin might be a deeply conserved feature of laminar growth, present even in a monolayered organ type.

Together, our findings position callose-dependent cytokinesis as a determinant of in-plane growth coordination that, beyond its role in cell division, safeguards against growth conflicts to maintain robust planarity in a monolayered tissue. By isolating this within-layer contribution from the inter-layer mechanics at play in angiosperm models, our study further establishes the moss phyllid as a minimal system for dissecting the general principles of planar organ morphogenesis in land plants (20).

## Material and methods

### Plant material and growth conditions

*Physcomitrium patens* Gransden ecotype was used as the genetic background for all experiments. Plants were grown *in vitro* under sterile conditions on BCD medium supplemented with dibasic ammonium tartrate (BCDAT medium) and solidified with 0.7% agar, following (50). Plants were maintained in a growth chamber at 23°C under a 16 h light/8 h dark photoperiod or continuous light for live-imaging experiments, with a light intensity of 50–80 μmol m⁻² s⁻¹. For live imaging, a transgenic line expressing *PpEF1α::acyl-YFP* (21) was used as the wild-type (WT) background for generating the callose synthase mutant *Ppcals6*.

### Generation and genotyping of *Ppcals6* mutants

To generate the *Ppcals6* knockout mutants, four protospacers were designed to target the exons 1, 3, 21 and 50 of the *PpCALS6* gene (Pp3c4_1790v3.1) using CRISPOR software (51). For enhanced readability, *PpCALS6* is referred to as *CALS6* throughout the manuscript. The corresponding sgRNAs (sgR-PpCALS6-EX01-59-F, CCATCCTGGGAACCGCACGCCTGG; sgR-PpCALS6-EX01-59-R, AAACCCAGGCGTGCGGTTCCCAGG; sgR-PpCALS6-EX03-33-F, CCATCCGTGAGAAGTCTGACATCT; sgR-PpCALS6-EX03-33-R, AAACAGATGTCAGACTTCTCACGG; sgR-PpCALS6-EX21-47-F, CCATGTACACTGTAGCATCGGCAT; sgR-PpCALS6-EX21-47-R, AAACATGCCGATGCTACAGTGTAC; sgR-PpCALS6-EX50-53-F, CCATTGCCGCTTGAAACAGGCGCA; sgR-PpCALS6-EX50-53-R, AAACTGCGCCTGTTTCAAGCGGCA) were synthesized, annealed and cloned into the destination vector pZeo-Cas9-gate published in (52), yielding the DM12-PpCALS6 plasmid. This construct was transformed into moss protoplasts of either wild type or the *PpEF1α::acyl-YFP* line using PEG-mediated method previously described (53). The transgenic lines were transiently selected based on antibiotic resistance and genotyped by PCR using the primers PD40_F6 (GGTGATTTCTCGAGGAGGGC) and PD40_R6 (CATACGGGCAATCTCTCGCA). Mutant lines with complete gene deletions were retained (lines #2 and #7 in the wild-type background, and lines #15, #19, #21 and #24 in the *PpEF1α::acyl-YFP* background), and all presented similar morphological defects. The lines #2 and #15 were used for experiments shown in Figure 1 and Figures 2-4, respectively. Predicted off-target sites suggested by CRISPOR software were screened by PCR, confirming the absence of non-specific mutations.

### Gene identifiers and expression data

Expression profiles of the *CALLOSE SYNTHASE* (*CALS*) gene family members were obtained using the expression dataset “*P. patens* vegetative development” of the PEATmoss database (54, 55) (https://peatmoss.plantcode.cup.uni-freiburg.de/easy_gdb/tools/expression/expression_input.php). *PpCALS4* corresponds to Pp3c10_19330V3.1 gene model, *PpCALS5* corresponds to Pp3c9_4140V3.1 gene model, *PpCALS6* corresponds to Pp3c4_1790V3.1 gene model, *PpCALS11* corresponds to Pp3c15_22960V3.1 gene model and *PpCALS12* corresponds to Pp3c4_15060V3.1 gene model. Throughout the manuscript, and unless otherwise stated, gene names are abbreviated for readability, *e.g. CALS6* refers to the *P. patens* gene *PpCALS6*.

### Microscopy

#### Gross phenotyping

Colony morphology, gametophore shoots, and individual phyllid outlines were imaged using a Keyence VHX-1000E digital microscope equipped with a 20–200× objective.

#### Live and fixed imaging of phyllids

Confocal images of individual phyllids were acquired using a Zeiss LSM 800 microscope equipped with a 40× water-immersion Apochromat objective (NA 1.0). Z-stacks were acquired at 0.5-μm intervals along the z-axis, with an image resolution of 1024 × 1024 pixels. YFP fluorescence from the membrane marker was excited using a 488-nm diode laser, and emission was collected over the 400–614-nm range. To image phyllids stained with propidium iodide (PI), the plant tissue was first immersed in a 50 μg/mL PI solution and subsequently scanned using 488 and 561 nm diode lasers. YFP and PI emissions were collected at 400–550 nm and 588–617 nm, respectively. For time-lapse imaging of upper phyllid development, gametophores bearing newly emerged 12th phyllids were selected from 18-day-old plates. Neighboring phyllids were carefully removed using fine tweezers to expose the targeted phyllid. The same phyllid was imaged at 12-h intervals over approximately six days, until maturity. When necessary, gametophores were tilted to expose the abaxial surface of the target phyllid. For samples extending beyond the objective’s field of view, multiple overlapping z-stacks were acquired to cover the entire phyllid. During imaging, plant samples were immersed in sterile water. After each imaging session, samples were rinsed three times with sterile water, gently blotted with tissue paper, and returned to their original growth conditions in the growth chamber. At least three independent time-lapse series were acquired for each experiment. Fine Z-stacks were acquired using DAPI imaging settings (excitation at 358 nm; emission at 461 nm) to visualize cell wall autofluorescence. Raw image data were processed using ZEN Black software (Carl Zeiss, Jena, Germany). Three-dimensional reconstructions in ZEN Blue were used to determine the number and spatial distribution of gaps.

#### Callose imaging

Aniline blue fluorochrome (Biosupplies) was prepared at 0.1 mg/mL in sterile deionized water and stored at 4 °C in the dark, according to (56). Before use, the stock was diluted 1:3 (33 µg/mL final) in 0.1 M phosphate buffer, pH 8.5 (97 mM K₂HPO₄, 3 mM KH₂PO₄). Gametophore apices/filaments were harvested from the edge of well-grown colonies, avoiding agar, and placed in 100 µL of staining solution per well in a 24-well plate. The plate was wrapped in aluminum foil and incubated for 30 min at room temperature. Samples were then mounted in distilled water between a slide and a coverslip. Imaging was performed on a Leica SP8 confocal microscope equipped with a 405 nm diode laser (10% power), a PMT spectral detector, and 10× dry and 40× water-immersion objectives. Aniline blue fluorescence was collected with a 440–520 nm emission band. The signal at the cell wall between the apical and subapical cells was first located under UV illumination with the 10× objective, then imaged with the 40× objective. Acquisition parameters were: frame size 1024 × 1024, 12-bit depth, zoom 1.5×, pinhole 1 A.U., 8-frame averaging, 8000 Hz scan speed. Z-stacks comprised 30 steps of 0.7 µm. Images were processed in Fiji.

### Image processing and quantitative analyses

#### Phyllid reconstruction

Confocal time-lapse images were processed using MorphoGraphX (57, 58). Images were subjected to two successive Gaussian blur operations, followed by Edge Detect using a threshold of 6,000–13,000 and Edge Detect Angle using a threshold of 4,000–8,000. Two-and-a-half-dimensional (2.5D) meshes were generated using a cube size of 5 μm, then subdivided and smoothed two to three times. Fluorescence signals were projected onto the meshes from a distance of 4–6 μm from the surface. The meshes were manually segmented, and parent cells were identified and labeled across consecutive imaging sessions. Cell lineages were reconstructed across multiple days based on parent-cell labels, and cell clone identities were assigned manually.

#### Cellular growth and division

Cellular growth and division were quantified using cell lineage information and the topology of neighboring cells. Area expansion was calculated from the total area of the daughter cells at time point (T_{i+1}) relative to the area of their parent cell at time point (T_i). Cell division was quantified as the number of daughter cells identified at the next time point minus one. The principal directions of growth between consecutive time points were characterized using the mutually perpendicular Stretch Max and Stretch Min measurements in MorphoGraphX.

#### Cell position

To quantify cell position along the phyllid, a group of cells at the phyllid base was manually labeled to define the basal boundary, which was positioned perpendicular to the main phyllid axis. The distance of each cell from this boundary was calculated using the Cell Distance function in MorphoGraphX, with Wall Weights set to Euclidean to obtain absolute distances. Relative distances from the phyllid base were calculated by normalizing absolute distances to the maximum distance measured at the corresponding time point. Distances from the phyllid tip were calculated using the same approach, with the single apical cell used as the reference.

#### Growth defects

To investigate the local effects of growth defects in *cals6* phyllids, growth-arrested or slow-growing cells were identified from heat maps of cell area expansion calculated over consecutive 12-h intervals. Direct neighboring cells sharing a common cell wall with the affected cells were also identified, excluding cells belonging to the midrib lineage. Cumulative area expansion was calculated for the affected cells and their neighbors from the onset of the growth defect until the phyllid first reached its mature shape. A corresponding group of cells in WT phyllids was analyzed for comparison.

#### Curvature

Cellular curvature was quantified using segmented meshes in MorphoGraphX. Mean cell curvature was calculated using the Project Mesh Curvature function and visualized as a heat map using the Heat Map – Measures – Signal Total function with default parameters.

#### Gametophore and phyllid length

The length of gametophores and the length, width, and perimeter of individual phyllids were measured using Fiji software (59). Shoot length was measured by tracing the gametophore stem using the segmented-line tool. Phyllid length was measured along the midrib, width was measured at the widest point, and perimeter was determined by tracing the phyllid contour using the segmented-line tool.

#### Cell wall gap size and thickness

Cell wall autofluorescence was used to examine the number, size, and distribution of cell wall gaps in *cals6* phyllids and to measure cell wall thickness in both wild-type (WT) and *cals6* phyllids. Phyllids were cleared for 14 days in a solution containing 120 g chloral hydrate, 7.5 mL glycerol, and 150 mL water. Cleared samples were embedded in the same solution and imaged using a Zeiss LSM 900 confocal microscope equipped with an Airyscan 2 detector and a C-Apochromat 40×/1.20 W objective, using multiplex Airyscan mode. Cell wall thickness and gap size were measured from two-dimensional images using ZEN Blue, with both transverse and longitudinal cell walls included in the analysis. Cell wall thickness was measured using the cell wall autofluorescence between two adjacent cell walls in at least 200 cells. Gap size in *cals6* was quantified from 350 measurements, and gap numbers were determined in more than 100 cells.

### Computational modeling

The model was implemented in the MorphoMechanX framework (60, 61), following the approach previously described in Lin et al. (21), as a 2D representation of the phyllid at two scales: cells, each refined into triangular finite elements that never cross cell boundaries. Growth parameters were assigned to each cell according to its type and position, and inherited by its elements. When a cell divided, its elements were discarded and the daughter cells were re-triangulated. Because residual stresses are released at each growth step, this remeshing did not alter the force distribution in the tissue. Growth was simulated with a growing finite element method (FEM), using 2D three-node triangular membrane elements instead of 3D six-node wedge elements, and a linear St. Venant isotropic material (Young’s modulus 100 MPa, Poisson’s ratio 0.3). At each step, the reference configuration of each element was enlarged, with independent control of growth parallel (K_par) and perpendicular (K_per) to a polarity direction, itself derived from the gradient of a distance field computed from the apical cell. Mechanical equilibrium was then solved, vertex positions updated and residual stresses released. To define the initial template, a representative day-1 mesh was flattened into 2D and its cell outlines extracted as polygons to serve as the template. The triangulation was updated upon division, with edge points regularized to maintain suitable triangle aspect ratios. The apical cell was given a distinct identity, and the most basal vertices were fixed along the longitudinal axis (Dirichlet conditions). Concerning tissue zonation and cell division, the phyllid was divided into three zones: a slow-growing basal attachment zone; above it, a proliferative zone defined by distance from the base (in cells), where cells divided on reaching a threshold area along the shorter of two candidate walls, oriented parallel or perpendicular to their polarity; and a differentiation zone, entered as cells were displaced out of the proliferative zone, where division stopped. The apical cell instead divided at 60° in an alternating left–right pattern. Growth rates could be set as gradients within zones, and zone sizes varied to match developmental phases and/or the influence of auxin. Zones and parameters were updated after each growth step. Growth arrest was simulated by specifying “wooden cells” that stop growing and dividing, and are 100-fold stiffer than other cells; these are shown in grey in the model output visualizations. At a defined time point, a midrib was specified from cells close to the base and to the central longitudinal axis. These cells received distinct growth parameters, notably a strong reduction of medio-lateral growth, and their identity was passed on to daughter cells.

## Acknowledgments

This work was supported by Czech Science Foundation grant 22-35680M (to R.P.) and 26-23380N (to P.C.), the Centre SÈVE, the Fonds de Recherche du Québec Nature et Technologies New Researcher grant 2020-NC-267497, and the Natural Sciences and Engineering Research Council of Canada Discovery grant RGPIN-2025-04418 (to D.K.), and a Biotechnological and Biological Sciences Research Council Institute Strategic Program Grant to the John Innes Centre BB/X01102X/1 (to R.S.S). We acknowledge the Imaging Facility of the IEB CAS, supported by MEYS CR LM2023050 ‘Czech-BioImaging’ and IEB CAS. Confocal imaging was mainly conducted using the instrument supported by Canada Foundation for Innovation grant (37805) (to D.K.).

## Author contributions

Conceptualization: W.L., D.K., and Y.C. ; Methodology: W.L., P.C., R.S.S., D.K., and Y.C. ; Software: M.D., D.G., R.S.S. ; Formal analysis: W.L., M.D., D.G., R.S.S., D.K., and Y.C. ; Investigation: W.L., S.H., P.C., M.D., D.G., A.L., and Y.C. ; Writing – original draft: W.L., D.K., and Y.C. ; Writing – review & editing: W.L., P.C., R.P., R.S.S., D.K., and Y.C. ; Visualization: W.L., D.K., and Y.C. ; Supervision: R.P., R.S.S., D.K., and Y.C. ; Project administration: D.K., and Y.C. ; Funding acquisition: R.P., D.K., and Y.C.

---

**Figure S1.**
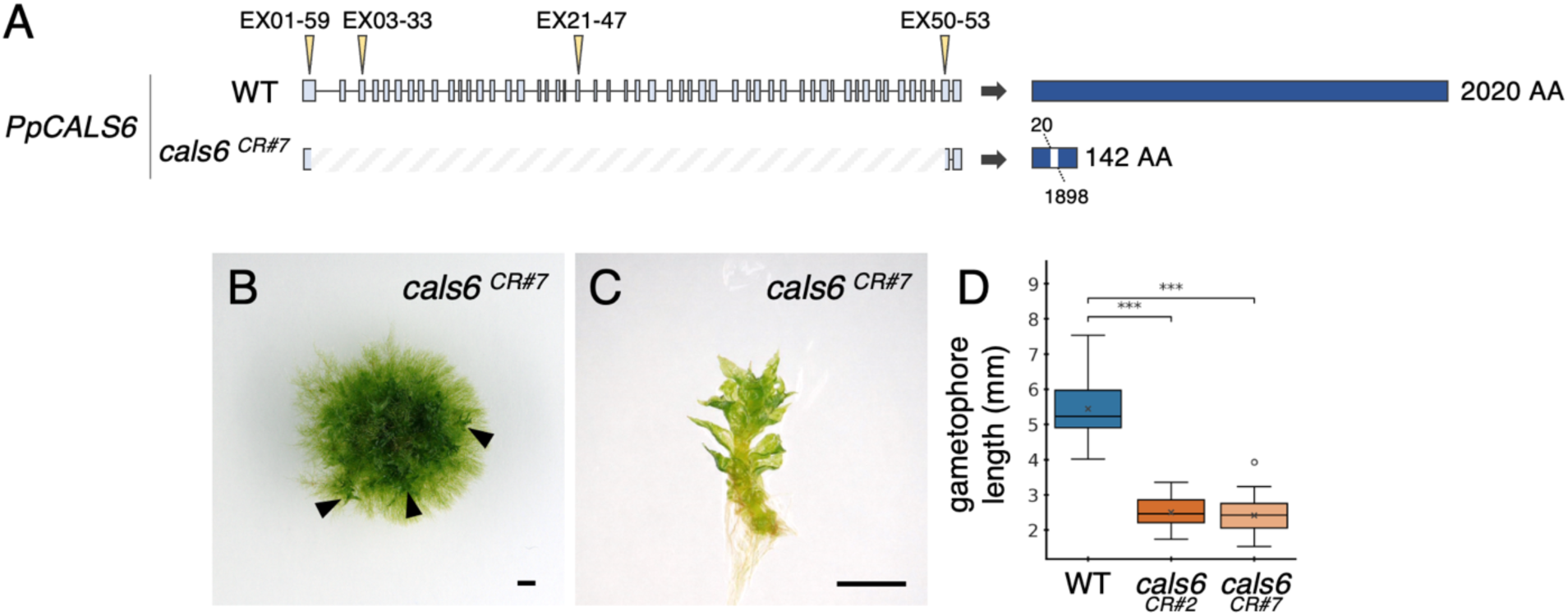
Molecular and phenotypic characterization of *cals6* CRISPR mutants. (**A**) Molecular characterization of the *cals6* CRISPR mutant line #7 (CR#7). Left: diagrams showing the *CALS6* locus in wild type (WT) and *cals6* mutant backgrounds, with light grey-blue boxes showing exons, yellow arrowheads indicate the positions of sgRNAs, and dashed lines indicate the edited region identified in the *cals6* mutant by Sanger sequencing. Right: schematic representations of the predicted proteins, with WT sequence shown in dark blue and predicted gap in the *cals6* mutant shown in white. Small numbers indicate predicted amino acid (AA) position or total number. (**B-C**) Representative colony (B) and gametophore (C) showing developmental alterations in *cals6^CR#7^* mutants (compare with WT in Figure 1). (**D**) Quantification of gametophore length (n = 20) in *cals6* mutants (line #2 refers to *cals6* mutant line shown in Figure 1), revealing that they are shorter than in WT. Statistical significance of a Welch t-test indicated as ***, (p ≤ 0.001). Scale bars : B-C, 1 mm.

